# ASFVdb: An integrative resource for genomics and proteomics analyses of African swine fever

**DOI:** 10.1101/670109

**Authors:** Zhenglin Zhu, Geng Meng

## Abstract

The recent outbreaks of African swine fever (ASF) in China and Europe have threatened the swine industry globally. To control the transmission of the African swine fever virus (ASFV), we developed ASFVdb, the African swine fever virus database, an online data visualization and analysis platform for comparative genomics and proteomics. On the basis of known ASFV genes, ASFVdb reannotates the genomes of every strain and annotates 4833 possible ORFs. Moreover, ASFVdb performs a thorough analysis of the population genetics of all the published genomes of ASFV strains and performs functional and structural predictions for all genes. For each ASFV gene, visitors can obtain not only basic information of the gene but also the distribution of the gene in strains, conserved or high mutation regions, possible subcellular location of the gene and topology of the gene. In the genome browser, ASFVdb provides sliding window population genetics analysis results, which facilitate genetics and evolutional analyses at the genomic level. The web interface is constructed based on SWAV 1.0. ASFVdb is freely accessible at http://asfvdb.popgenetics.net.

## Introduction

African swine fever (ASF), caused by the ASF virus (ASFV), is a viral disease in domestic or wild swine (Parker et al., 1969; Thomson et al., 1980; Anderson et al., 1998). The mortality rate of ASF is nearly 100% for infected domestic swine. The outbreaks in China and central Europe in the summer of 2018 caused important economic losses in the global swine industry. However, efforts to produce effective vaccines or treatments for ASFV are obstructed by the complexity of the virus. The ASF virus, which has an icosahedral capsid with a multilayered membrane structure, contains many polypeptides, the function and identities of which are largely unknown. It has been reported that some generated antibodies are not protective but may enhance the disease (Detray, 1963; Stone and Hess, 1967; Mebus, 1988; Blome et al., 2014; Sunwoo et al., 2019) due to the complexity of the virion and the intracellular and extracellular localization of infectious particles. Consequently, identification of functional ASFV polypeptides is important to develop effective vaccines.

Due to the above observations, we developed ASFVdb, the African swine fever database, to assist ASFV research. Different from previous relevant ASFV databases, such as ViralZone (Hulo et al., 2011), ASFVdb is specific for ASFV. ASFVdb compares all the published ASFV genomes (Chapman et al., 2008; de Villiers et al., 2010; Chapman et al., 2011; Bishop et al., 2015; Portugal et al., 2015; Olesen et al., 2018; Zani et al., 2018; Bao et al., 2019), annotates each genome and validates the persistence of each gene in different strains. ASFVdb was developed to facilitate the identification of gene function and identity. Aside from a basic annotation, it also provides information on the subcellular location of genes and functions of genes, topology predictions, comparative genomic alignments and population genetics analysis results.

## Result and discussions

### Overview of the genomic data in ASFVdb

ASFVdb collects a broad range of published ASFV genome data (Table S1) and analyzes and identifies 253 ASFV gene clusters (Table S2) from the genomes of 38 strains (for details, see Material and Methods). Among the 253 gene clusters, 145 have conserved ORFs in all strains (Table S2). Reannotation of the genomes of the 38 strains predicted 4833 genes and 924 genetic remains (Figure 1). In each strain, there are 181-263 genes, on average. These results did not contradict previous reports of 151-181 genes per strain (Galindo and Alonso, 2017; Bao et al., 2019), considering that the pipeline included all possible alternative splices (Figure S1) and the discrepancies between annotations for different strains with different strategies at different times (Chapman et al., 2008; de Villiers et al., 2010; Chapman et al., 2011; Bishop et al., 2015; Portugal et al., 2015; Olesen et al., 2018; Zani et al., 2018; Bao et al., 2019). We performed a subcellular localization analysis of the ASFV genes to determine their roles in the infection process (Figure 2A). We found that more than half of the ASFV (60%) are single-pass or multi-pass membrane proteins (Figure 2B). 8% of the ASV genes (23 cases) are capsid proteins, which may provide potential targets for vaccine design. In gene ontology, the largest number of ASFV genes are enriched in the membrane and had catalytic activity (Figure 3). Moreover, based on the population genetics test analysis, we found that 334 genes have a significantly low Tajima’s D (Tajima, 1989) and significantly high composite likelihood ratio (CLR) (Nielsen et al., 2005; Zhu and Bustamante, 2005) (Rank Test, P-value<0.05), indicating that these genes were recently possibly under positive selection (Table S3). 77 genes have a significantly high Tajima’s D value (Table S4, Rank Test, P-value<0.05), and they may be involved in balancing selection. These results are informative for future ASFV research.

**Figure 1.**
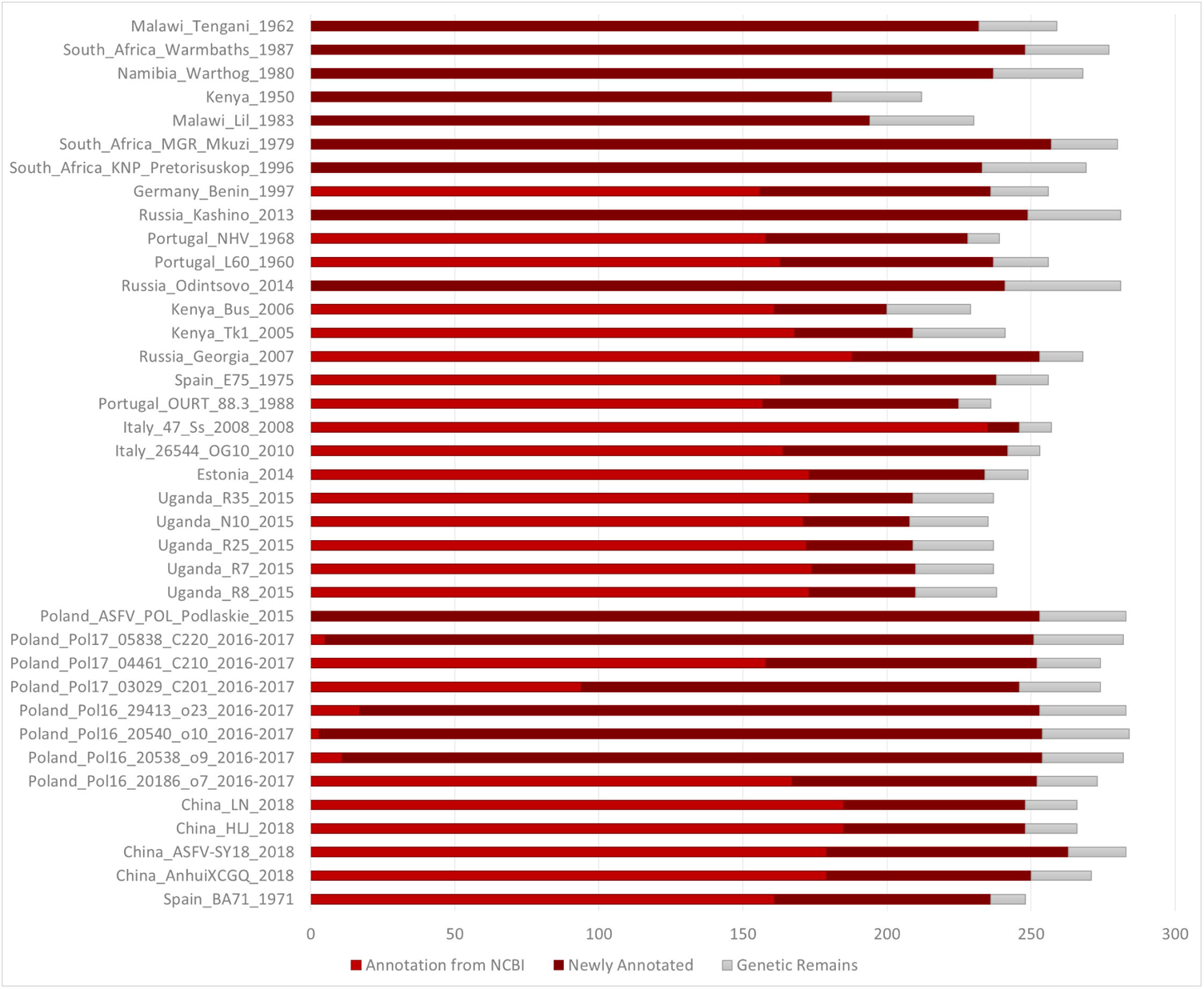
Number of annotated genes in all the ASFV strains. Genes annotated in NCBI, genes newly annotated and genetic remains are marked in dark red, red and gray, respectively.

**Figure 2.**
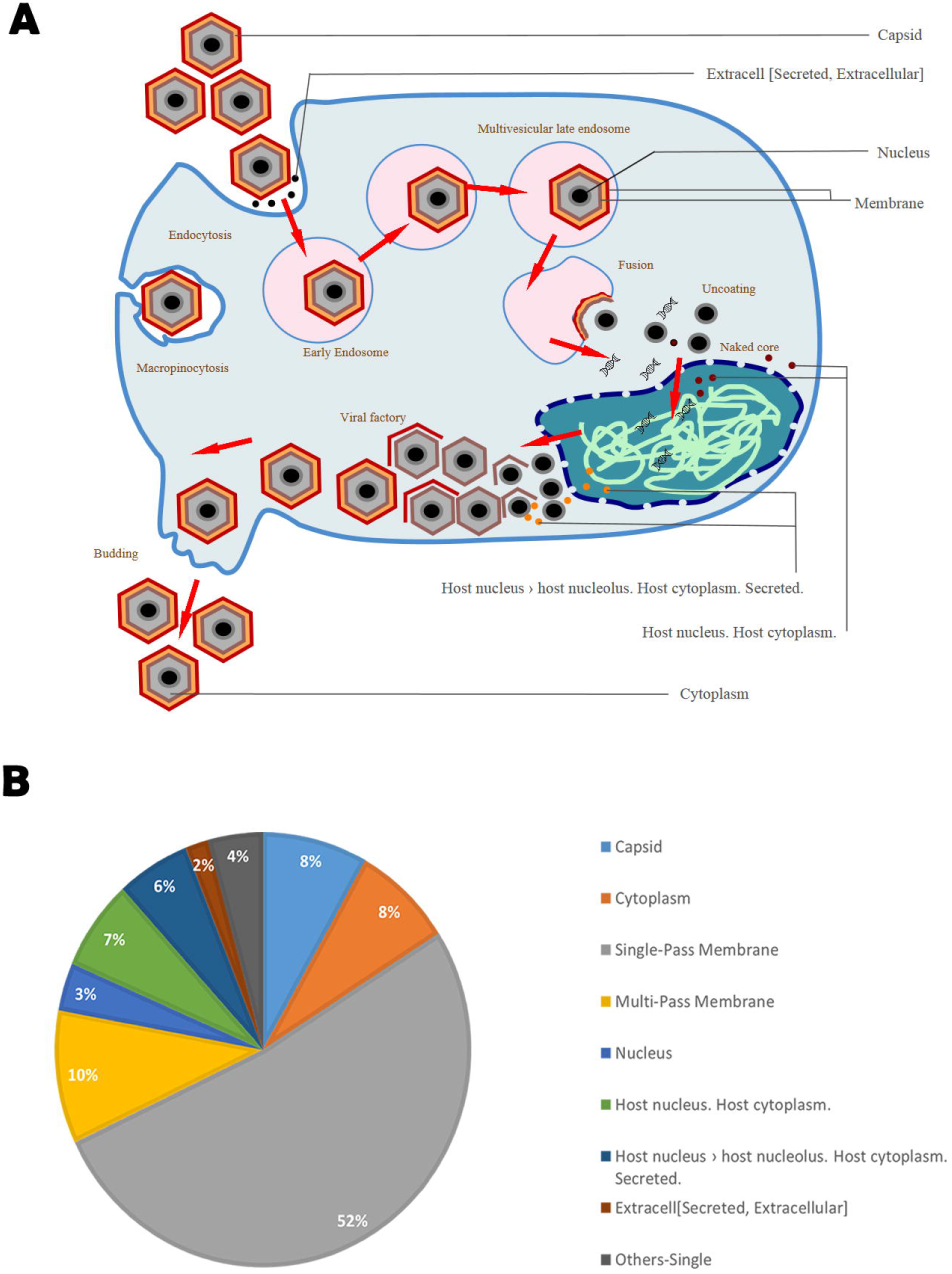
A shows a diagram of the infection process of ASFV with the MSLVP subcellular locations marked. B shows the percentages of ASFV gene clusters in each subcellular location group.

**Figure 3.**
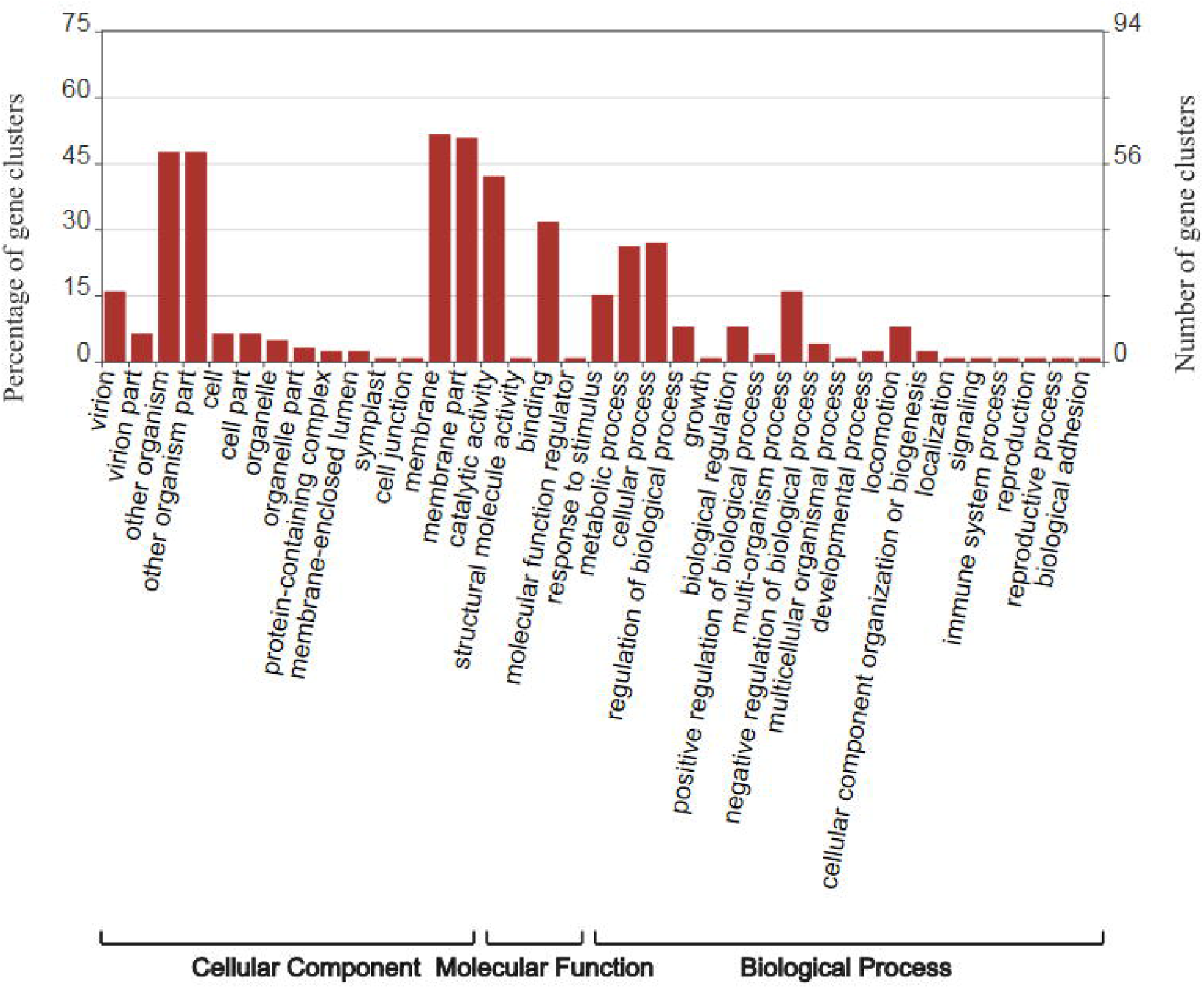
Enrichment of the gene ontology (GO) of ASFV gene clusters. This figure was drawn by WEGO (Ye et al., 2006).

### ASFVdb offers a diverse search method

The main search input of ASFVdb, located at the center of the home page, allows users to search a gene by its general gene name, gene accession or description of the gene’s function. We created a gene alias list to make the search results as complete as possible. In the right column of the home page, there is a list of other gene search methods in ASFVdb. First, users can BLAT against one ASFV genome, like the workflow in the UCSC genome browser (Karolchik et al., 2007), or BLAST against CDS (coding sequence) or protein sequences. Second, users can search for genes in a specific subcellular location. In the results view, we list the topology information and function annotation of each gene to allow users to conveniently localize the gene in the host-virus interaction. Third, users can find genes that have a specific function according to the gene’s GO annotation. “Gene Clusters” lists the persistence of the ASFV genes in the 38 strains. Users can view a specific gene by clicking an item in the gene list table. Moreover, to more conveniently track a list of genes, ASFVdb provides gene links by inputting a list of the genomic positions or a list of the accession numbers of genes. These operations facilitate analysis of ASFVdb with a personal gene list.

### Evolutionary episodes of the ASFV strains in ASFVdb

On the home page, below the main search box, a phylogenetic tree displays the evolutionary history of the available published 38 ASFV strains. In contrast to the previous method used to make ASFV phylogenetic trees (de Villiers et al., 2010; Bao et al., 2019), we built the tree by performing multiple alignments on the genomic scale to better understand the processes and patterns of evolution (Wilgenbusch et al., 2017). The tree we constructed (Figure 4) is consistent with previous publications (Galindo and Alonso, 2017; Bao et al., 2019; Sanchez et al., 2019). Our tree reflects transmission events during the spread of ASFV. After it was first identified in Kenya in the 1920s (Montgomery, 1921), ASFV was subsequently reported in eastern and southern Africa (Thomson G. R., 2004). In the mid-last century, ASFV spread from Africa to Portugal (Sanchez-Botija, 1963) as its first stop in Europe. Then, ASFV spread to Malta (1978), Italy (1967, 1980), France (1964, 1967, 1977), Belgium (1985) and the Netherlands in 1986 (Costard et al., 2009). In 2007, ASFV was introduced to Caucasus (Rowlands et al., 2008) and began to spread across Russia and Eastern Europe. In 2018, ASFV was found in China, and outbreaks in China and central Europe enhance the seriousness of this disease (OIE, 2019).

**Figure 4.**
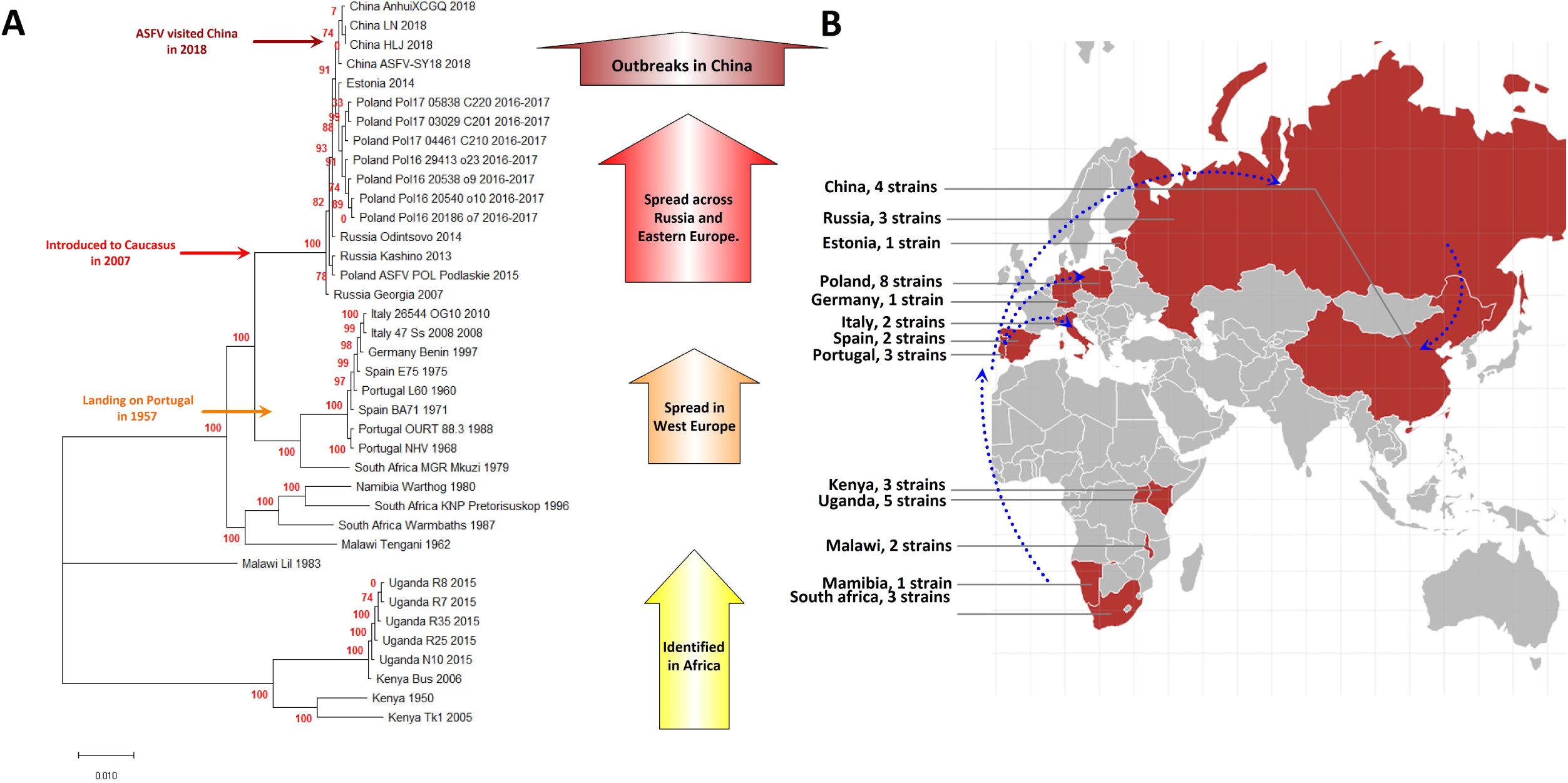
A, Phylogenetic tree of the ASFV strains in ASFVdb with historical events marked. The numbers marked in red are the marginal likelihood of the tree. B. The distribution of strains in ASFVdb according to country. The blue arrows with dotted lines indicate the possible infection paths of ASFV.

### Gene Visualization in ASFVdb

Through clicking the taxon name in the tree or selecting an item in the “Gbrwoser” list, users can go to the genome browser page (Figure 5), where gene segments are subsequently arranged along the genome according to the genomic positions of genes. Genes annotated from the NCBI GFF file are colored in blood red as “Annotation from NCBI”. Genes annotated by mapping (for details, see the Material and Methods) are colored in dark red as “Newly annotated”. “Genetic Remains” are colored in light gray. The gene name is provided above each gene segment, and “>” or “<“ indicates a gene’s transcriptional direction. Four tracks of population genetics analysis, Pi, Theta (Watterson, 1975), Tajima’s D (Tajima, 1989), and CLR (Nielsen et al., 2005; Zhu and Bustamante, 2005) are listed following the gene track. To help trace selective signatures, we drew a top 5% line for CLR and two 5% lines in ascending and descending order for Pi, Theta and Tajima’s D. Using the tool bar at the top, users can move or zoom the genome in the browser, set the focus bar to a selected region, export the drawing of one track with gene segments and export the data of one track.

**Figure 5.**
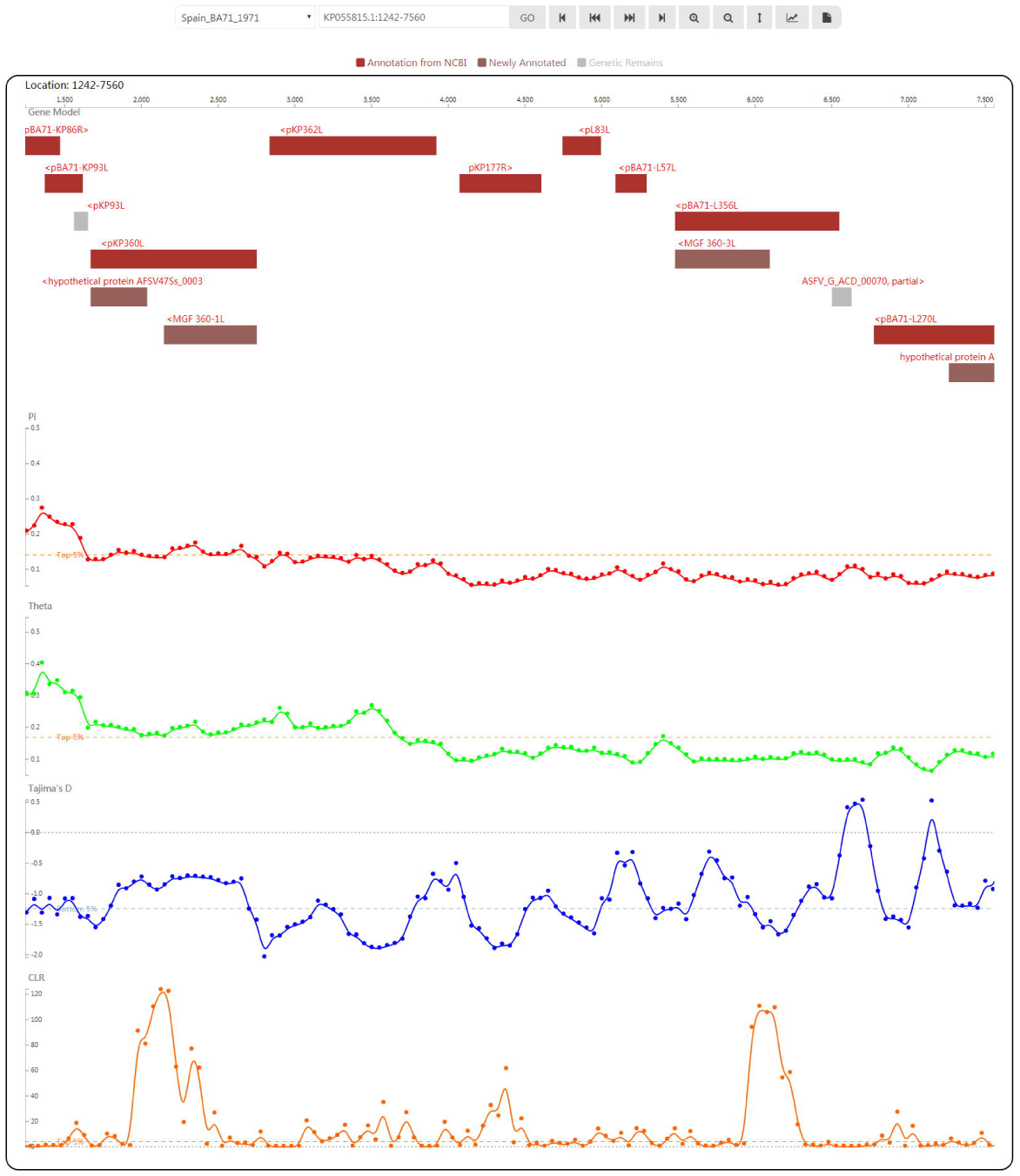
Gnome browser view to display ASFV genes and population genetics test statistics.

Clicking one gene segment will lead to the gene’s information page. The full list of annotations includes basic information (strain, gene name, description, location in the genome, Genbank Accession and full name), sequence (CDS and protein), summary (function, UniProt accession, related Pubmed ID, related EMBL ID, corresponding Proteomes ID, related Pfam ID and correlated Interpro ID), ontologies (GO and KEGG), subcellular location, topology (transmembrane region prediction), genomic alignment in the CDS region, multiple alignment of orthologues, gene tree of corresponding NCBI annotated or newly annotated proteins, and orthologous genes in strains. Texts that link to internal or external sites are hyperlinked to facilitate viewing and analysis.

### Analysis of CD2v (EP402R) in ASFVdb: a case study

CD2v (EP402R) is an ASFV gene related to hemadsorption (Rodriguez et al., 1993). CD2v (EP402R) plays a key role in viral attenuation and may be involved in the generation of immune protection from the host against ASFV infection (Monteagudo et al., 2017). Through searching for “EP402R” in the strain China_AnhuiXCGQ_2018, we found that this gene encodes a single-Pass membrane protein in the transmembrane region from 207 AA to 229 AA (Figure S2A). In the genome view with the population genetics analysis tracks, we found that the transmembrane region is composed of a Tajima’s D valley and a CLR peak (Figure S2B), inferring the selection signatures. The region from 1AA to 206AA is predicted to be located outside of the viron membrane, with peaks of Pi and Theta, indicating its high diversity among strains. From the information regarding “Genomic alignment in the CDS region” and “Orthologues in Strains”, we found that this gene is only highly conserved among 27 of all 38 compared strains (Figure S2C). This result may partly explain why pigs vaccinated with baculovirus-expressed EP402R/CD2v did not have full protection from ASFV infection (Ruiz-Gonzalvo et al., 1996; Escribano et al., 2013). AFSVdb annotates that this protein has similarity (E-value=0.000801919) to the cell adhesion molecule CD2 (Jones et al., 1992), the crystal structure of which (Figure S2D, from PDB) was used to build a homology modelling structure of EP402R.

## Conclusion

To assist veterinary scientists and researchers to combat the threat of ASFV outbreaks, we developed a database-based toolbox, ASFVdb. ASFVdb integrates data from NCBI, UniProt (Apweiler et al., 2004), ViralZone (Hulo et al., 2011) and a broad range of published articles (Hulo et al., 2011). ASFVdb specializes in ASFV. ASFVdb compares all published ASFV genomes (Chapman et al., 2008; de Villiers et al., 2010; Chapman et al., 2011; Bishop et al., 2015; Portugal et al., 2015; Olesen et al., 2018; Zani et al., 2018; Bao et al., 2019), summarizes data on the proteins of all known ASFV strains, and provides not only basic information of genes but also external links, subcellular localization, topology, comparative genomic alignments, a gene tree and evolutionary analysis results. This information is helpful and convenient for the further accession of proteomics and biochemical data in ASFV research. Hopefully, ASFVdb will be useful for further research on ASFV and vaccine design.

## Materials and methods

### Data collection and processing

We downloaded the genome sequences, CDS, proteins and relevant annotations of 38 sequenced ASFV strains (Table S1) from the NCBI genome database. Among the 3 different versions (NC_001659.2, U18466.2 and KP055815.1) of Spain_BA71_1971 genome data, we used the original version, KP055815.1. Of the 38 different ASFV genomic data, only 23 are fully annotated, and their gene IDs or names are not unified. To harmonize this inconsistency, we constructed a unified dataset and used the dataset to reannotate the 38 genomes. Specifically, we performed pairwise alignments for all of the annotated protein sequences. The sequences were clustered together with an identity > 50% and coverage > 80% according to previous homologous gene identification methods (She et al., 2009; Yue et al., 2018). Finally, a list containing 308 unified genes was generated. Then, we mapped these unified genes to all 38 genomes. If a unique protein mapped with the highest score of identity > 50% and coverage > 80%, we considered the protein to be a unique protein in the strain, and the mapped region in the strain genome was marked. To identify the corresponding mapped region of one unique gene that might express a protein, we translated the mapped region using NCBI ORFfinder and aligned all the possible translations of the unique protein. We took the alignment with the highest score and checked whether its identity and coverage were both higher than 0.5. If its identity and coverage were both higher than 0.5, we consider the region to express proteins; if not, we marked the region as “Genetic Remains”. The genes that were annotated in the NCBI genome database were marked as “Annotation from NCBI”, and unannotated gene regions with a high expression probability were marked as “Newly Annotated”. Finally, all of the sequenced genes were clustered into 253 gene clusters with CD-HIT (Li and Godzik, 2006) and required identification > 50% and coverage > 70%.

To retrieve the annotation of the ASFV gene product from UniProt (Apweiler et al., 2004; Patient et al., 2008), we BLASTed the protein sequence of the ASFV gene against the UniProt protein database (Patient et al., 2008) and took the hits with the highest scores as the best matches. If the best match of one ASFV gene had an E-value <0.05, we extracted the accession number of the mapped sequence as the UniProt ID of the matched ASFV gene. With the UniProt ID, we obtained the external annotations of the ASFV gene, such as the corresponding PubMed ID, EMBL ID, Proteomes ID, Pfam, Interpro, Gene Ontology, and KEGG and function annotations. We also BLASTed all of the ASFV protein sequences against the PDB database (Berman et al., 2000; Burley et al., 2019) in the same way and recorded the alignment results. Disappointedly, we only obtained 45 unique proteins from the PDB that had an E-value <0.05, which means that most of the ASFV-expressed proteins did not have structural information.

### Subcellular localization and topology prediction

The subcellular location and topology annotations from UniProt only cover several ASFV genes. Thus, we reperformed these predictions for all of the genes. We predicted the subcellular location of the ASFV proteins through MSLVP (Thakur et al., 2016) using the parameters of “one-versus-one”, “Second-tier Algorithm” and a similarity >90%. A total of 253 gene clusters were grouped into 9 subcellular locations. We predicted transmembrane helixes within the protein sequences using TMHMM 2.0 (Krogh et al., 2001). The output images were converted into PNG format to be displayed on the website by Magick (www.imagemagick.org).

### Comparative genomics and population genetics analysis

To trace the evolutionary history of ASFV, we performed a whole genome alignment of all 38 ASFV genomes by Mugsy (Angiuoli and Salzberg, 2011) and built a phylogenetic tree by FastTree 2.1 (Price et al., 2010), with the parameter ‘-boot 5000’ to test the likelihood of the generated tree. To obtain a more detailed comparative genomic map of the 38 strains, we used LASTZ (Harris, 2007) to perform genome-genome alignments between any two ASFV strains and outputted the results in AXT format. For one ASFV gene, we retrieved the corresponding sequences of other strains from the genome-genome alignment results, realigned these sequences by MUSCLE (Edgar, 2004b; Edgar, 2004a), and obtained the genomic alignment of the CDS region. According to the persistence of each gene in different strains, we also retrieved all of the protein counterparts of one unique gene and performed a multiple alignment by MUSCLE. To more clearly display the evolutionary history of each protein, we also drew trees from the multiple alignment as we did for the genome.

We used a window of 200 bp and a step size of 50 bp to slide along the ASFV genome. From the genome-genome alignments described above, we retrieved sequences within the sliding window that we used to calculated Pi, Theta (Watterson, 1975) and Tajima’s D (Tajima, 1989) using VariScan 2.0 (Vilella et al., 2005; Hutter et al., 2006). We wrote Perl scripts to call the allele frequencies and used SweapFineder2 (DeGiorgio et al., 2016) to calculate the composite likelihood ratio (CLR) (Nielsen et al., 2005; Zhu and Bustamante, 2005) with a step size of 50. We took the median of the population genetics test statistics in the region of each gene as the corresponding value.

### Web Interfaces

The web interface of ASFVdb was developed using Mysql + PHP + CodeIgniter (www.codeigniter.com) + JQuery (jquery.com) and was based on SWAV (swav.popgenetics.org), which is an open source web application for population genetics visualization and analysis. We used (Yachdav et al., 2016) to show the multiple alignments of orthologous CDS or proteins and phylotree (Shank et al., 2018) to display the phylogenetic trees of genomes or proteins. We made changes to the source code of the web application to fit the development of ASFVdb, such as adding links within the diagram. The search interfaces of the sequence alignments were constructed based on PHP parsing of the results from BLAT (Kent, 2002) and NCBI BLAST (Johnson et al., 2008). The workflow of the other search blocks, such as searching by the molecular structure or gene names, are mostly SQL query pipelines.

## Data availability

All ASFVdb data are publicly and freely accessible at http://asfvdb.popgenetics.net. Feedback on any aspect of the ASFVdb database and discussions of ASFV gene annotations are welcome by email to.

## Acknowledgments

This work was supported by grants from the National Natural Science Foundation of China (31200941) and the Fundamental Research Funds for the Central Universities (106112016CDJXY290002).

