## Supplementary Figures for "ASFVdb: An integrative resource for genomics and proteomics analyses of African swine fever"

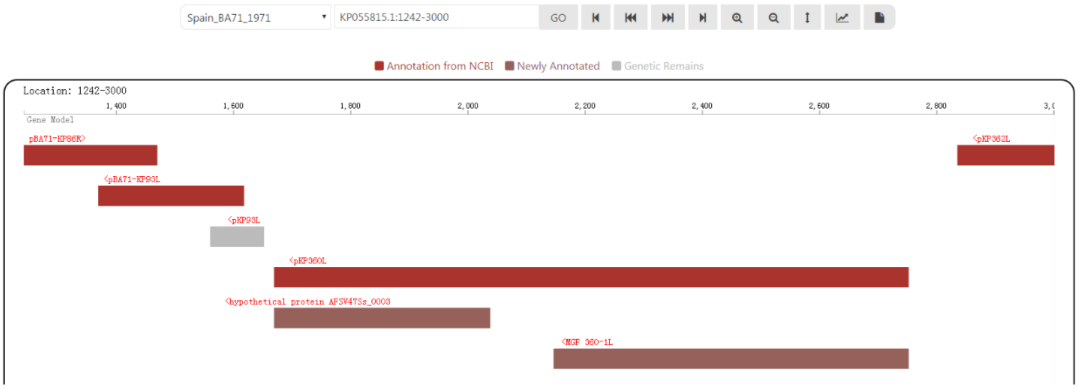


Figure S1. A snapshot displaying the inclusion of alternative splicing cases in ASFVdb. As shown is the figure, 'pKP360L' colored in red is of possibility to be transcribed in the forms of "hypothetical protein AFSV47Ss_0003" and "MGF 360-1L", colored in terracotta.

**A**

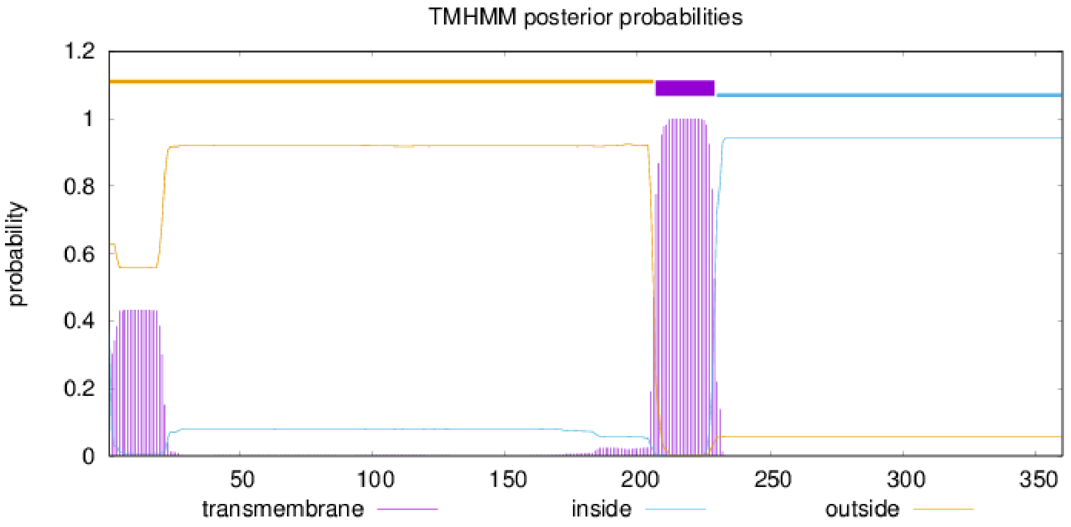
TMHMM posterior probabilities
probability
1.2
1
0.8
0.6
0.4
0.2
0
50
100
150
200
250
300
350
transmembrane
inside
outside

# B

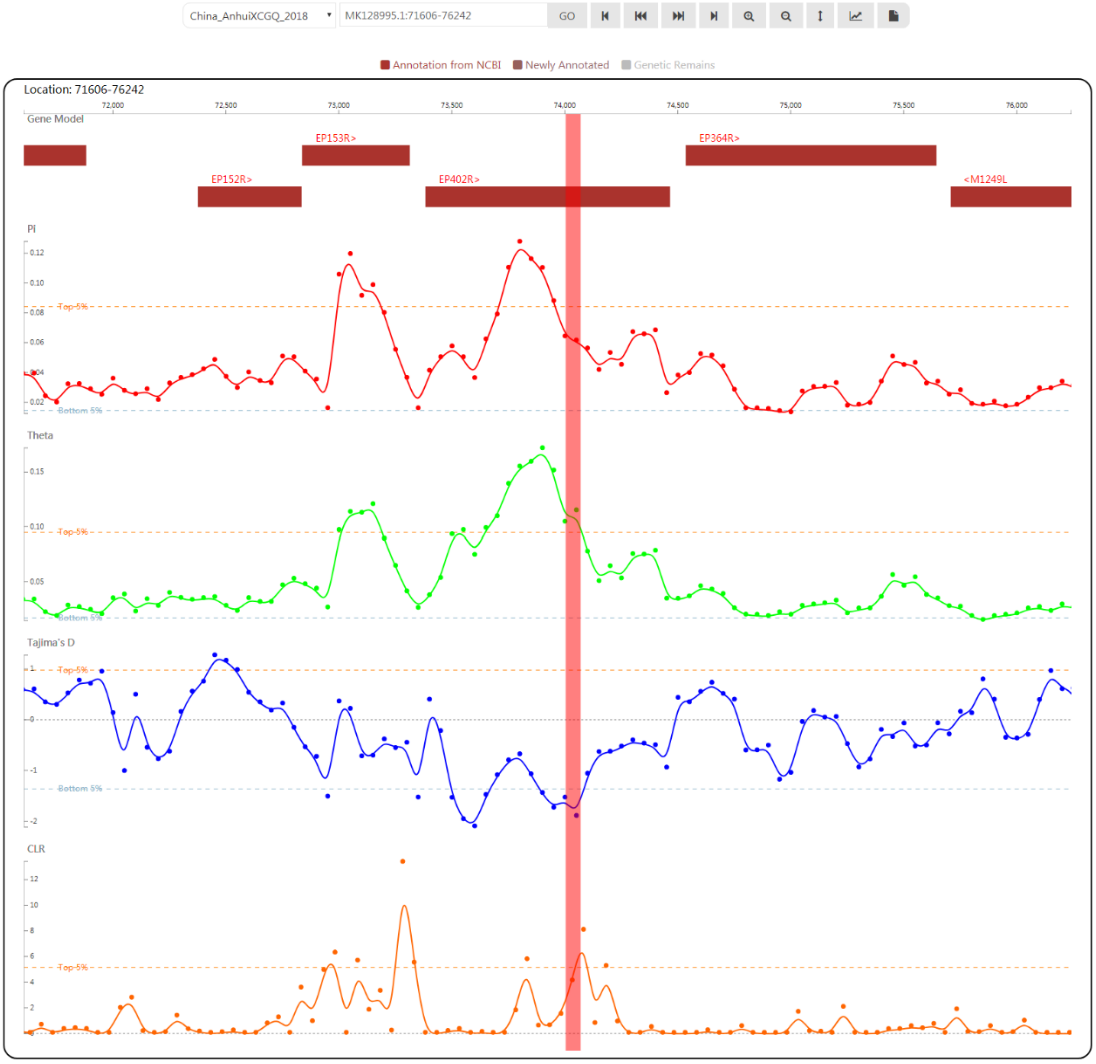

China_AnhuiXCGQ_2018
MK128995.1:71606-76242
GO
Annotation from NCBI
Newly Annotated
Genetic Remains
Location: 71606-76242
72,000
72,500
73,000
73,500
74,000
74,500
75,000
75,500
76,000
Gene Model
EP153R>
EP364R>
EP152R>
EP402R>
<M1249L
Pi
Theta
Tajima's D
CLR
Top-5%
Bottom 5%

# C

Orthologous in Strains

| Strain | Availability | Status | Gene |
|---|---|---|---|
| Spain_BA71_1971 | ✔ | Protein | AKO62740.1 |
| China_AnhuiXCGQ_2018 | ✔ | Protein | AYW34030.1 |
| China_ASFV-SY18_2018 | ✔ | Protein | CBW46724.1 |
| Poland_Pol16_20186_o7_2016-2017 | ✔ | Protein | AXZ95833.1 |
| Poland_Pol16_20538_o9_2016-2017 | ✔ | Protein | CBW46724.1 |
| Poland_Pol16_20540_o10_2016-2017 | ✔ | Protein | CBW46724.1 |
| Poland_Pol16_29413_o23_2016-2017 | ✔ | Protein | CBW46724.1 |
| Poland_Pol17_03029_C201_2016-2017 | ✔ | Protein | AXZ96021.1 |
| Poland_Pol17_04461_C210_2016-2017 | ✔ | Protein | AXZ96116.1 |
| Poland_Pol17_05838_C220_2016-2017 | ✔ | Protein | CBW46724.1 |
| Poland_ASFV_POL_Podlaskie_2015 | ✔ | Protein | CBW46724.1 |
| Uganda_R8_2015 | ✖ | | |
| Uganda_R7_2015 | ✖ | | |
| Uganda_R25_2015 | ✖ | | |
| Uganda_N10_2015 | ✖ | | |
| Uganda_R35_2015 | ✖ | | |
| Estonia_2014 | ✔ | Protein | SPS73481.1 |
| Italy_26544_OG10_2010 | ✔ | Protein | AJZ77076.1 |
| Italy_47_Ss_2008_2008 | ✔ | Protein<br>Protein | AKO62740.1<br>AOO54390.1 |
| Portugal_OURT_88.3_1988 | ✔ | Protein | CAN10406.1 |
| Spain_E75_1975 | ✔ | Protein | CBH29159.1 |
| Russia_Georgia_2007 | ✔ | Protein | CBW46724.1 |
| Kenya_Tk1_2005 | ✔ | Protein | AJL34072.1 |
| Kenya_Bus_2006 | ✖ | | |
| Russia_Odintsovo_2014 | ✔ | Genetic Remain | CBW46724.1 |
| Portugal_L60_1960 | ✔ | Protein<br>Genetic Remain | AIY22249.1<br>CAN10406.1 |
| Portugal_NHV_1968 | ✔ | Protein | AIY22408.1 |
| Russia_Kashino_2013 | ✔ | Protein | CBW46724.1 |
| Germany_Benin_1997 | ✔ | Protein | CAN10158.1 |
| South_Africa_KNP_Pretorisuskop_1996 | ✖ | | |
| South_Africa_MGR_Mkuzi_1979 | ✖ | | |
| Malawi_Lil_1983 | ✔ | Genetic Remain | CBW46724.1 |
| Kenya_1950 | ✖ | | |
| Namibia_Warthog_1980 | ✔ | Genetic Remain | CBW46724.1 |
| South_Africa_Warmbaths_1987 | ✖ | | |
| Malawi_Tengani_1962 | ✖ | | |
| China_HLJ_2018 | ✔ | Protein | QBH90546.1 |
| China_LN_2018 | ✔ | Protein | QBH90731.1 |

**D**

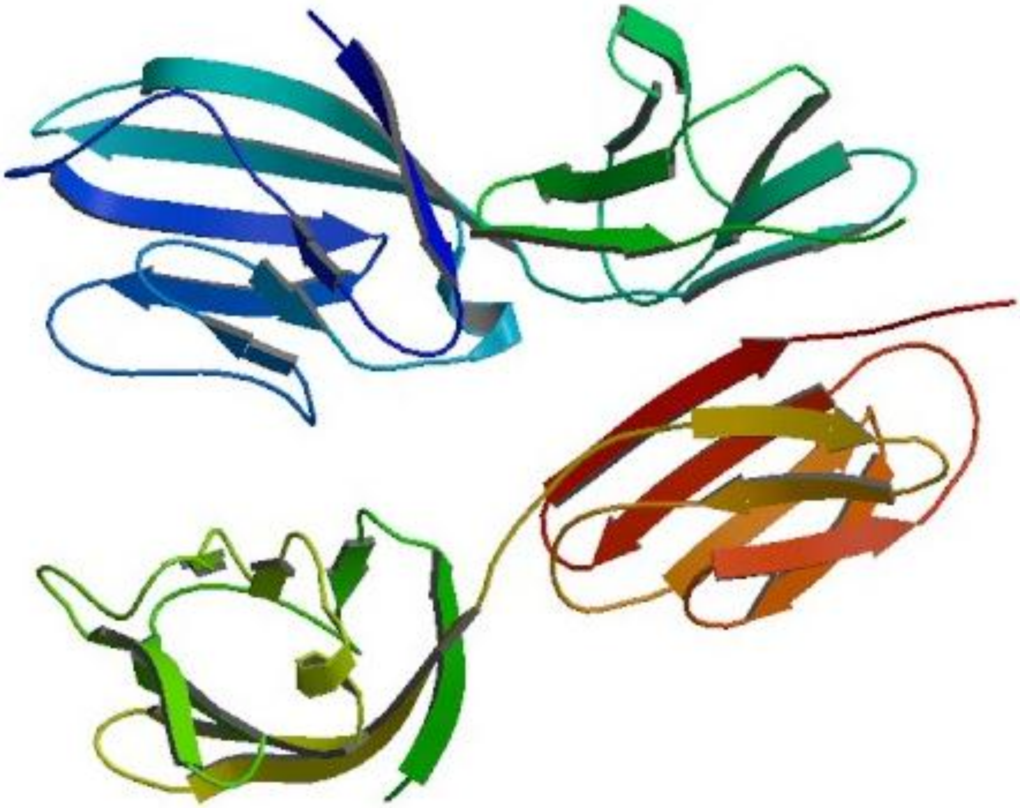

Figure S2.

Analysis results of EP402R by ASFVdb. A is the transmembrane prediction of EP402R in China_AnhuiXCGQ_2018. B is the population statistics test results in China_AnhuiXCGQ_2018, where red bar marked the transmembrane region of EP402R. C is the persistence of EP402R in different strains. D is the 3D structure of the cell adhesion molecule CD2 from PDB, which is of similarity to EP402R (E-value<0.05).
